# Enantioselectivity in the Enzymatic Dehydration of Malate and Tartrate: Mirror Image Specificities of Structurally Similar Dehydratases

**DOI:** 10.1101/2023.06.17.545066

**Authors:** Asutosh Bellur, Souradip Mukherjee, Pragya Sharma, Vijay Jayaraman, Hemalatha Balaram

**Author notes:** School of Biology, Indian Institute of Science Education and Research Thiruvananthapuram, Maruthamala Campus, Vithura, Kerala 695551, India (Vijay Jayaraman).

## Abstract

Malate (2-hydroxysuccinic acid) and tartrate (2,3-dihydroxysuccinic acid) are chiral substrates; the former existing in two enantiomeric forms (R and S) while the latter exists as three stereoisomers (R,R; S,S; and R,S). Dehydration by stereospecific hydrogen abstraction and *anti-* elimination of the hydroxyl group yield the achiral products fumarate and oxaloacetate, respectively. Class-I fumarate hydratase (FH) and L-tartrate dehydratase (L-TTD) are two highly conserved enzymes belonging to the iron-sulfur cluster hydrolyase family of enzymes that catalyze reactions on specific stereoisomers of malate and tartrate. FH from *Methanocaldococcus jannaschii* accepts only S-malate and S,S-tartrate as substrates while the structurally similar L-TTD from *Escherichia coli* accepts only R-malate and R,R-tartrate as substrates. Phylogenetic analysis reveals a common evolutionary origin of L-TTDs and two-subunit archaeal FHs suggesting a divergence during evolution that may have led to the switch in substrate stereospecificity preference. Due to the high conservation of their sequences, a molecular basis for switch in stereospecificity is not evident from analysis of crystal structures of FH and predicted structure of L-TTD. The switch in enantiomer preference may be rationalised by invoking conformational plasticity of the amino acids interacting with the substrate, together with substrate reorientation and conformer selection about the C2-C3 bond of the dicarboxylic acid substrates. Although classical models of enzyme-substrate binding are insufficient to explain such a phenomenon, the enantiomer superposition model suggests that a minor reorientation in the active site residues could lead to the switch in substrate stereospecificity.

## Introduction

Enantioselectivity in enzymatic reactions involving chiral substrates is an almost universal feature in biochemistry. Most enzymes are exclusively selective catalysts that catalyze reactions on only one enantiomer. There are varied ways by which enzymes have evolved to work around stereoisomers, with evolution of different protein folds with distinct active site architectures being one among them(Lamzin, Dauter, & Wilson, 1995; McCarter & Stephen Withers, 1994). Tartrate dehydratases eliminate water from tartrate, the molecule which in Pasteur’s hands led to the birth of modern stereochemistry(Pasteur, 1848). Class-I fumarate hydratase (FH) and L-tartrate dehydratase (L-TTD) are members of Fe-S cluster containing hydrolyase family of enzymes sharing a high sequence similarity but catalyzing reactions on substrates of opposite stereochemistry(Flint & Allen, 1996; Reaney, Begg, Bungard, & Guest, 1993). Although the primary reaction of FH is the interconversion of S-malate and fumarate, the enzyme also converts S,S-tartrate to oxaloacetate(Kronen, Sasikaran, & Berg, 2015; van Vugt-Lussenburg, van der Weel, Hagen, & Hagedoorn, 2009, 2013) whereas L-TTD catalyzes the stereospecific conversion of R-tartrate to oxaloacetate(Reaney et al., 1993). In malate or 2-hydroxybutanedioate (2-hydroxysuccinic acid), the C2 chiral carbon is either in S (L-malate) or R (D-malate) absolute configuration whereas in tartrate or 2,3-dihydroxybutanedioate (2,3-dihydroxysuccinic acid) both C2 and C3 carbons are chiral with (S, S)-tartrate (D-tartrate), (R, R)-tartrate (L-tartrate) and (2R, 3S)/2S, 3R)-tartrate (meso-tartrate) as possible diastereomers (Fig. 1C). Hereafter, for clarity, the stereoisomers of malate and tartrate will be referred to by R/S nomenclature only. All L-TTDs are made up of two subunits, α and β(Reaney et al., 1993). Class-I FHs from eukaryotes and many bacteria are single chain enzymes comprised of an N-terminal domain (NTD) and a C-terminal domain (CTD) while those from archaea are the two-subunit type with α and β subunits(Shimoyama, Rajashekhara, Ohmori, Kosaka, & Watanabe, 2007). The NTDs of single chain FH, the α subunits of archaeal FH, and the α subunits of L-TTD share high sequence similarity. Similarly, the β subunits of the two-subunit class-I FHs, and L-TTDs, and the CTDs of single chain FH possess high sequence similarity. Hence, using sequence information, L-TTD and class-I FH are hard to distinguish and have often been misannotated in the RCSB and UniProt databases.

**Figure 1.**
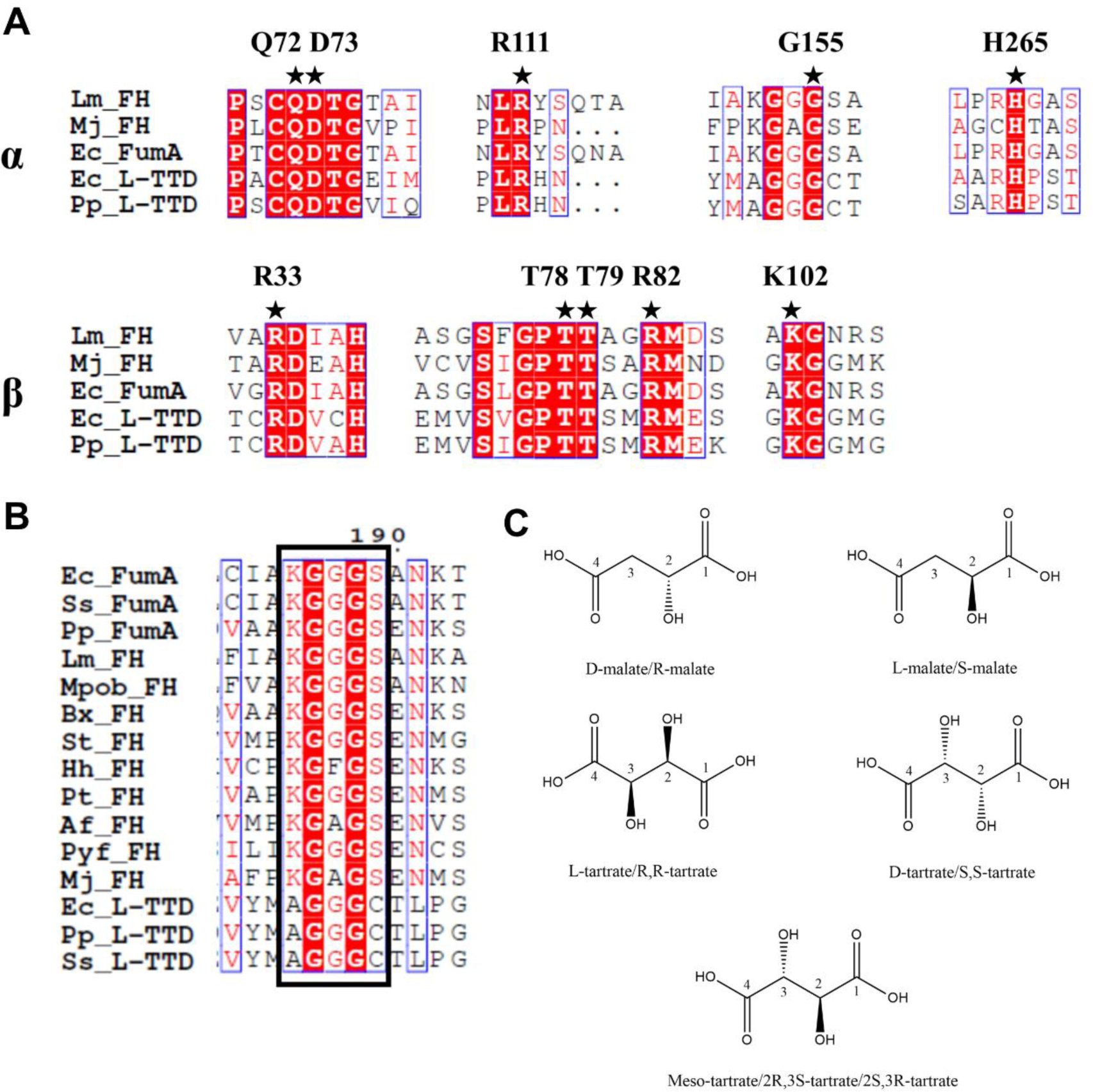
Active site and Proximal loop residues in class-I FHs and L-TTDs that utilize tartrate/malate of opposite stereospecificity. **A**. Multiple sequence alignment of class-I FHs and L-TTDs highlighting the active site residues involved in substrate binding and catalysis. Active site residues deciphered from the structure of *Lm*FH and *Mj*FH are indicated with a star. α, α subunit of *Mj*FH and L-TTD; β, β subunit of *Mj*FH and L-TTD. *Ec*L-TTD*, E. coli* L-TTD; *Pp*L-TTD, *Pseudomonas putida* L-TTD; Ec_FumA, *E. coli* fumarase A (FH); Lm_FH, *L. major* FH; Mj_FH, *M. jannaschii* FH. **B.** Multiple sequence alignment of KGXGS and AGGGC motifs of the Proximal loop present in biochemically characterised class-I FHs and L-TTDs, respectively. Complete sequence alignment and organism names are provided in Supplementary Figure S1. **C.** Structures of L/R,R-tartrate, D/S,S-tartrate, L/S-malate, D/R-malate, and meso/R,S/S,R-tartrate generated using ChemDraw. Tartrate is different from malate by the presence of an extra hydroxyl group on C3 carbon and two chiral centers, resulting in diastereomers.

Although preliminary characterization of L-TTD from *Pseudomonas putida*(Kelly & Scopes, 1986), *Escherichia coli*(Reaney et al., 1993) and an L-tartrate fermenting anaerobic bacteria(Schink, 1984) has been reported, no structure of L-TTD is available. Class-I FHs have been investigated extensively with crystal structures of the enzyme from *Leishmania major* (*Lm*)(Feliciano, Drennan, & Nonato, 2016) and *Methanocaldococcus jannaschii* (*Mj*)(Bellur et al., 2023) showing identical active site residues in similar conformation, the roles of which in substrate binding and catalysis have been confirmed by extensive mutagenesis. Due to the high sequence similarity, the modelled structure of L-TTD is similar to FH with identical active site architecture. This raises the issue of how the active sites of FH and L-TTD recognize tartrate of opposite stereochemistry. We have biochemically characterized *E. coli* (*Ec*) L-tartrate dehydratase. Various substrate analogues of tartrate have been screened for activity on both *Ec*L-TTD and *Mj*FH to understand the discrimination based on stereospecificity in these enzymes. Finally, a possible explanation for opposite substrate stereospecificity preference in these enzymes has been proposed using the enantiomer superposition model(Bearne, 2020; Bentley, 2003).

## Results and discussion

### 1. Molecular phylogeny of Class-I FH and L-TTD

Alignment of the protein sequences of biochemically characterized class-I FHs and L-TTDs from diverse organisms revealed high degree of conservation (Fig.S1). The active site residues that interact with S-malate in *Lm*FH are invariant across class-I FH and L-TTD sequences (Fig.1A). However, the alignment shows a key difference between L-TTDs and FHs in the sequence of the loop that is proximal to the active site in *Lm* and *Mj* FH crystal structures. This loop, referred to as Proximal Loop has a highly conserved KGXGS motif in FH (residues 144 to 148 in *Mj*FH α subunit), the sequence of which is AGGGC in L-TTD (residues 152 to 156 in *Ec*L-TTD) (Fig. 1B).

The two-subunit L-TTD is found only in bacteria while the two-subunit class-I FH is archaeal in origin. A phylogenetic tree created with a dataset of 264 FH and L-TTD sequences revealed two distinct clades (Fig. 2). The phylogenetic proximity of L-TTDs to two-subunit FHs suggests a possible horizontal gene transfer (HGT) between the two clades. This proximity between the two dehydratases has been observed earlier in an analysis carried out on a smaller number of sequences(Kronen & Berg, 2015). In addition, the phylogenetic tree shows that the single chain FHs from protozoa and actinobacteria form an independent clade while those from proteobacteria and firmicutes arise from a common node that contains L-TTDs and two-subunit FHs.

**Figure 2.**
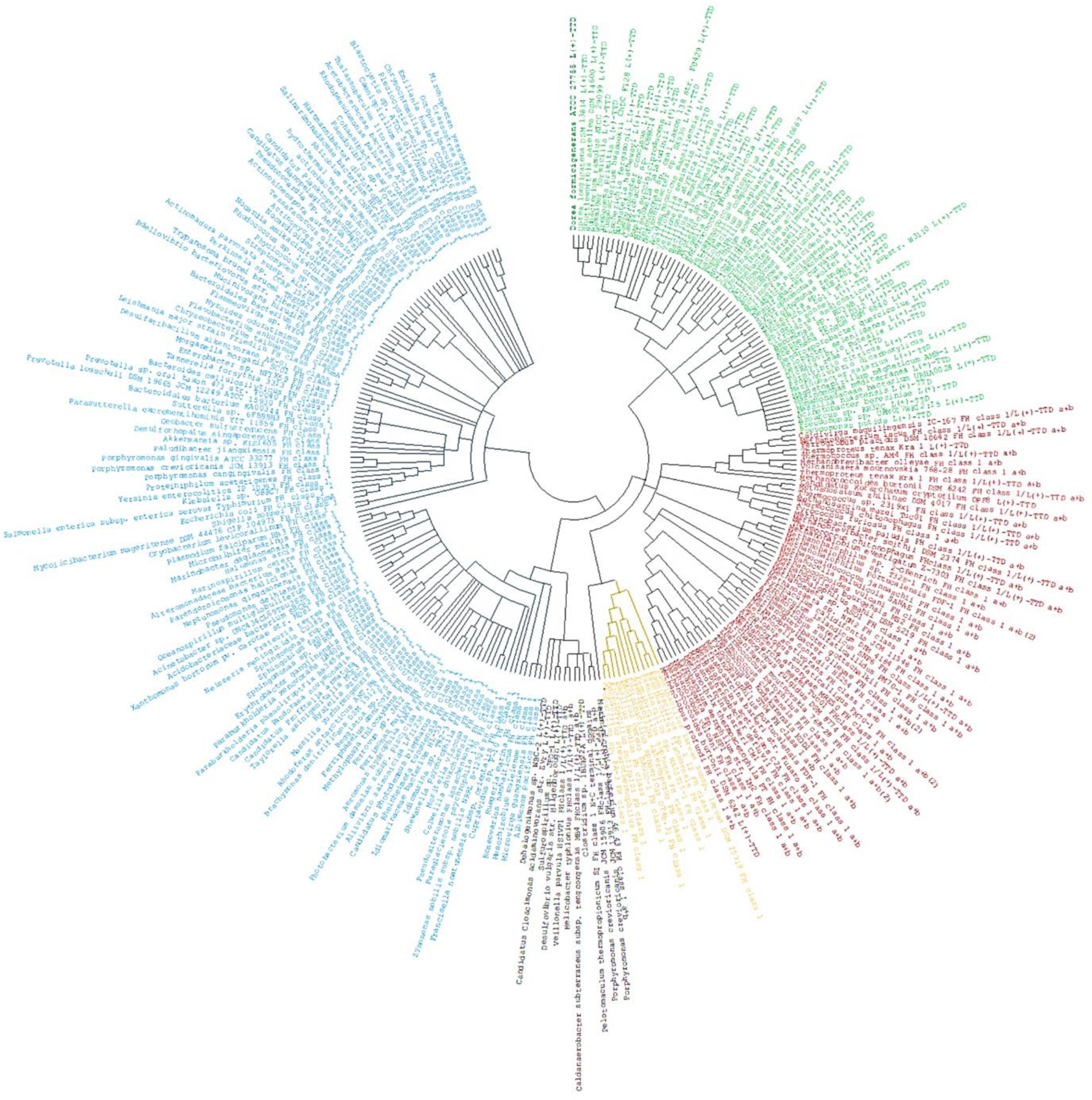
Phylogenetic tree (cladogram) of Class-I FH and L-TTD protein sequences. Single-subunit class-I FH and concatenated sequences of two-subunit class-I FH/L-TTD, representing all the taxa in which these proteins are present were used for the analysis. The tree was constructed using the Maximum Likelihood method and Le Gascuel model performing 500 bootstrap replicates(Felsenstein, 1985). Highlighted in blue are the single-subunit class-I fumarases; red, archaeal two-subunit class-I fumarases; green, bacterial L-tartrate dehydratases; yellow, single-subunit *Bacilli* class-I fumarases; and black, sequences annotated as either or both FH/L-TTD and having the KGXGS motif.

The high similarity between the sequences of the two classes of enzymes, complete conservation of active site residues suggest that L-TTD and FH have similar binding and catalytic mechanisms. It is interesting to note that all the L-TTD sequences used in the analysis have AGGGC sequence in the motif while all the FHs have the KGXGS with the exception of FHs from *Bacilli* that have KGGGC motif. *Bacilli* FH could be an intermediate in the HGT process from archaeal FH to bacterial L-TTD considering its origin from the same ancestral node.

### 2. Substrate specificity of *Mj*FH and *Ec*L-TTD

Apart from activity on S-malate (Bellur et al., 2023), as shown in Fig. S2A, *Mj*FH was found to be active on (S, S)-tartrate corroborating previous reports on other class-I FHs(Kronen & Berg, 2015). The enzyme was inactive on (R, R)-tartrate and R-malate. Testing the activity of *Mj*FH on the substrate S-malate in the presence of (R, R)-tartrate or R-malate, all at 1 mM concentration in the assay mixture, there was no reduction in the activity of the enzyme (Fig. S2B), going to show that these enantiomers/diastereomers of the substrates do not even bind to the enzyme’s active site. Like in other class-I FHs meso-tartrate was a potent competitive inhibitor of *Mj*FH, with a *K_i_* value of 161±17 μM. (Fig. S2C). The absence of inhibition by (R, R)-tartrate and R-malate indicates that *Mj*FH must bind 2S, 3R meso-tartrate where the active site is specific for the S configuration at C2 and relaxed at C3, binding both R and S configurations.

*Ec*L-TTD α and β subunits co-purified yielding a brown colored protein solution (Fig. 3A, B) that required pretreatment with (NH_4_)_2_Fe(SO_4_)_2_ and Na_2_S for obtaining active enzyme. This indicates that though the iron-sulfur cluster is bound to the purified enzyme, the required 4Fe-4S stoichiometry is achieved only after reconstitution. The molecular mass of the protein on size-exclusion chromatography corresponded to a dimer of a heterodimer (2α+2β subunits) with circular dichroism spectrum of the 4Fe-4S reconstituted enzyme exhibiting positive Cotton effect (Fig. S3A, B, C). Kinetic characterization yielded a *V*_max_ value of 12 ± 0.4 μmol min^-1^mg^-1^, *K_m_* value of 190 ± 30 μM, and *k_cat_*/*K_m_* value of 5.88 x 10^4^ s^-1^M^-1^ with (R, R)-tartrate as substrate (Fig. 3C). The enzyme was found to be moderately active on R-malate (1 mM) with specific activity of 0.2 μmol min^-1^ mg^-1^, which was 50 folds lower than for its substrate (R, R)-tartrate. S-malate, though found to be not a substrate, was a competitive inhibitor with a *K_i_* of 261 ± 29 μM (Fig. 3D). The enzyme was found to not bind (S,S)-tartrate as the activity remained unchanged in its presence. Similar to class-I FH, meso-tartrate was a strong competitive inhibitor of *Ec*L-TTD, inhibiting the enzyme with a *K_i_* of 149 ± 14 μM (Fig. 3E). Taken together, the ability of *Ec*L-TTD to catalyze the reaction on the substrates (R, R)-tartrate and R-malate, and be inhibited by S-malate, clearly indicates that both R or S configurations of C2 can bind to the active site whereas only the R configuration of C2 undergoes catalysis. In contrast, the C3 configuration has to be only R for both binding and catalysis. The also indicates that as in *Mj*FH, *Ec*L-TTD must bind 2S, 3R meso-tartrate.

**Figure 3.**
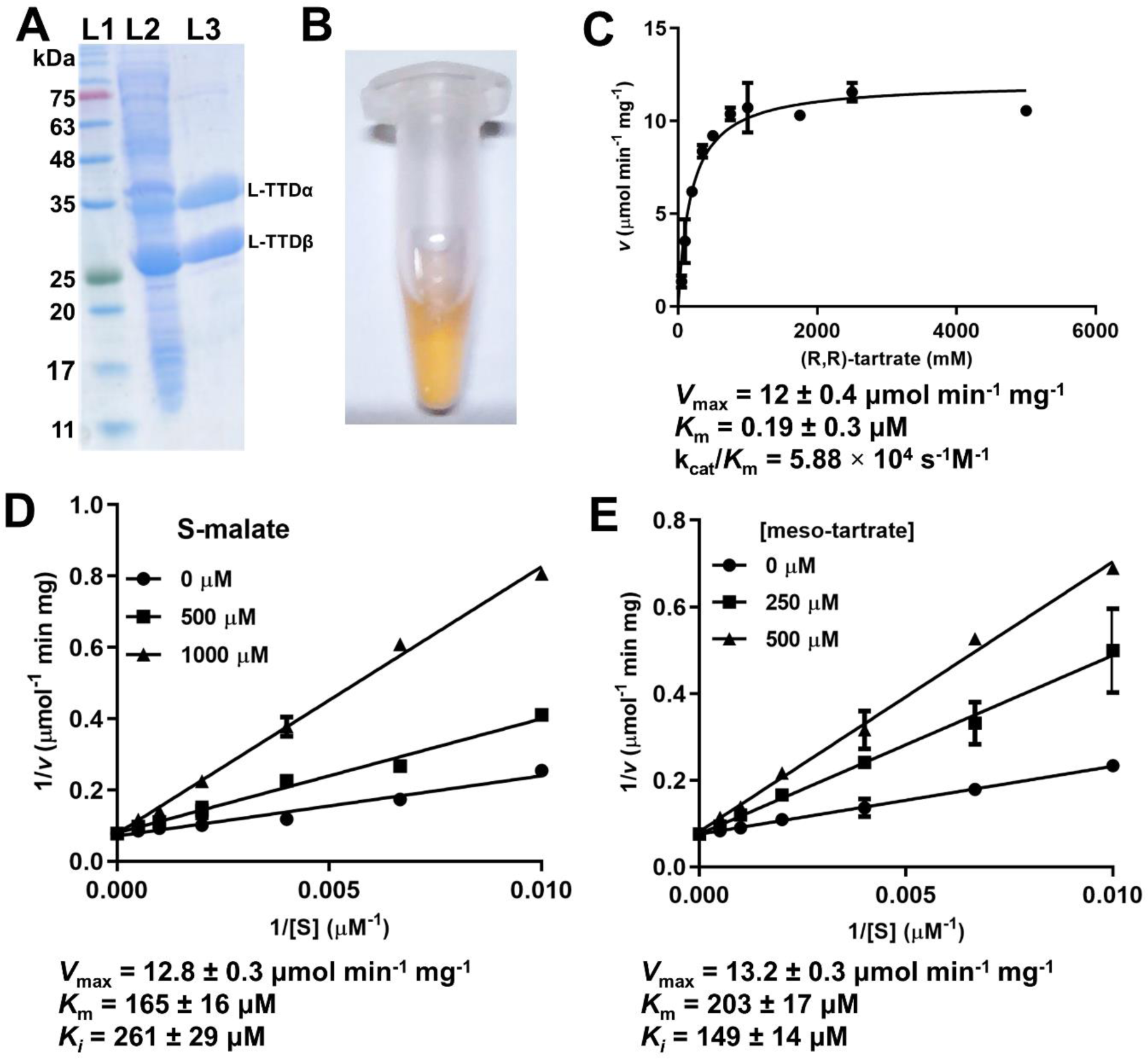
Kinetic characterization of *Ec*L-TTD. A. Purification of *Ec*L-TTD as monitored by SDS-PAGE. L1, protein molecular mass marker; L2, *E. coli* lysate expressing the subunits of L-TTD; L3, purified *Ec*L-TTD α (upper band) and β (lower band) subunits. **B.** Freshly purified *Ec*L-TTD shows brown coloration characteristic of iron-sulfur cluster proteins. **C.** Plot of initial rate, *v vs* R,R-tartrate concentration using reconstituted *Ec*L-TTD. Kinetic parameters obtained from fit to Michaelis-Menten equation are provided beneath the panel. **D.** Double reciprocal plots of 1/*v vs* 1/[S] with S-malate as inhibitor and R,R-tartrate as substrate. Concentration of R,R-tartrate was varied at different fixed concentrations of S-malate (0, 500 and 1000 μM). **E.** Double reciprocal plots of 1/*v vs* 1/[S] with meso-tartrate as inhibitor and R,R-tartrate as substrate. Concentration of R,R-tartrate was varied at different fixed concentrations of meso-tartrate (0, 250 and 500 μM). Assays in panel D and E were repeated twice to confirm reproducibility.

As the proximal loop motif (KGXGS/AGGGC) is a key difference between the two groups of enzymes, the motif in *Mj*FH and neighboring residues were replaced with those in *Ec*L-TTD (Fig. S4A, B). Overall, 4 mutants of *Mj*FH were generated, and activity and switch in substrate specificity were examined (Fig. S4C, D). All mutants showed no change in substrate stereospecificity preference, and only a decrease in activity on the substrate S-malate was observed. Further, inclusion of (R, R)-tartrate had no effect on activity indicating that this diastereomer does even bind to the mutants of *Mj*FH (Fig. S4E, F). The decrease in activity could be due to disruption of interactions of some of the residues in the loop with either the bound ligand or active site residues, noteworthy among them being the contacts between NH of G147 and C2-OH of malate, and between G145/A146 and the catalytic acid D62 (*Mj*FH numbering). This indicates that though the motif is proximal to the active site in class-I FH crystal structures and specific to the two classes of hydratases, it alone does not contribute to enantiomer/diastereomer specificity as mutations lead only to a decrease in activity.

### 3. Enantiomer superposition model addresses substrate stereospecificity

Our attempts to determine the crystal structure of *Ec*L-TTD were not successful. Multiple crystallization attempts with the unreconstituted freshly purified protein and with the Fe-S cluster reconstituted protein did not yield crystals. Attempts at obtaining the apoprotein failed and the protein remained colored despite treatment with EDTA and other chelating agents to remove the cluster. Hence, predicted structures were used for analysis of the active site architecture. The AlphaFold predicted structures of *Ec*L-TTD α-(AF-P05847-F1) and β-subunits (AF-P0AC35-F1) superposed well on *Mj*FH (RMSD_α_, 1.104 Å; RMSD_β_, 1.008 Å) and *Lm*FH (RMSD_α_, 1.95 Å; RMSD_β_, 1.52 Å) crystal structures (Fig. S5A, B). Moreover, the structure predicted by Baker and colleagues before any class-I FH structure was experimentally determined, shows overall similarity in folds indicating that the two groups of enzymes, being similar in sequence, have the same three-dimensional structure as well(Ovchinnikov et al., 2015). Examination of the AlphaFold predicted structure of *Ec*L-TTD α-subunit also revealed that the cysteine residues C71, C190, and C277, previously identified to co-ordinate with the cluster iron atoms (Bak & Weerapana, 2023), are well-positioned to accommodate the Fe-S cluster (Fig. S5C).

Examination of the crystal structure of *Lm*FH bound to S-malate shows that C4 carboxylate of the bound substrate is held by R173, R421 and Q134 while the C1 carboxylate is held by K491 and R471. The C3 carbon is in close proximity to the catalytic base T467 while C2OH contacts the catalytic acid D135 and the labile Fe in the cluster. This apart G216 of the KGXGS motif contacts C2OH (Fig. 4A). All these residues superpose well on the structure of *Mj*FH(Bellur et al., 2023). Examination of the active site in the modeled structure of *Ec*L-TTD did not reveal any difference in the orientation of the ligand-binding and catalytic residues, that could account for the altered substrate stereospecificity preference in L-TTD compared to class-I FHs (Fig. 4A, B). Hence, we resorted to the use of enantiomer superposition model to comprehend ligand binding and catalysis in *Ec*L-TTD(Bearne, 2020; Bentley, 2003). This model considers the three binding determinants (substituents) on the chiral carbon to lie on the same plane (pseudo-mirror plane), with flexibility in the binding of the 4^th^ substituent on the enantiomeric ligand, such that in either enantiomer it can interact with the same binding determinant on the enzyme. In this model, the 4^th^ binding determinant is oriented out of the plane at an angle of about 109° instead of pointing in the opposite direction as invoked in the earlier four location model(Mesecar & Koshland, 2000).

**Figure 4.**
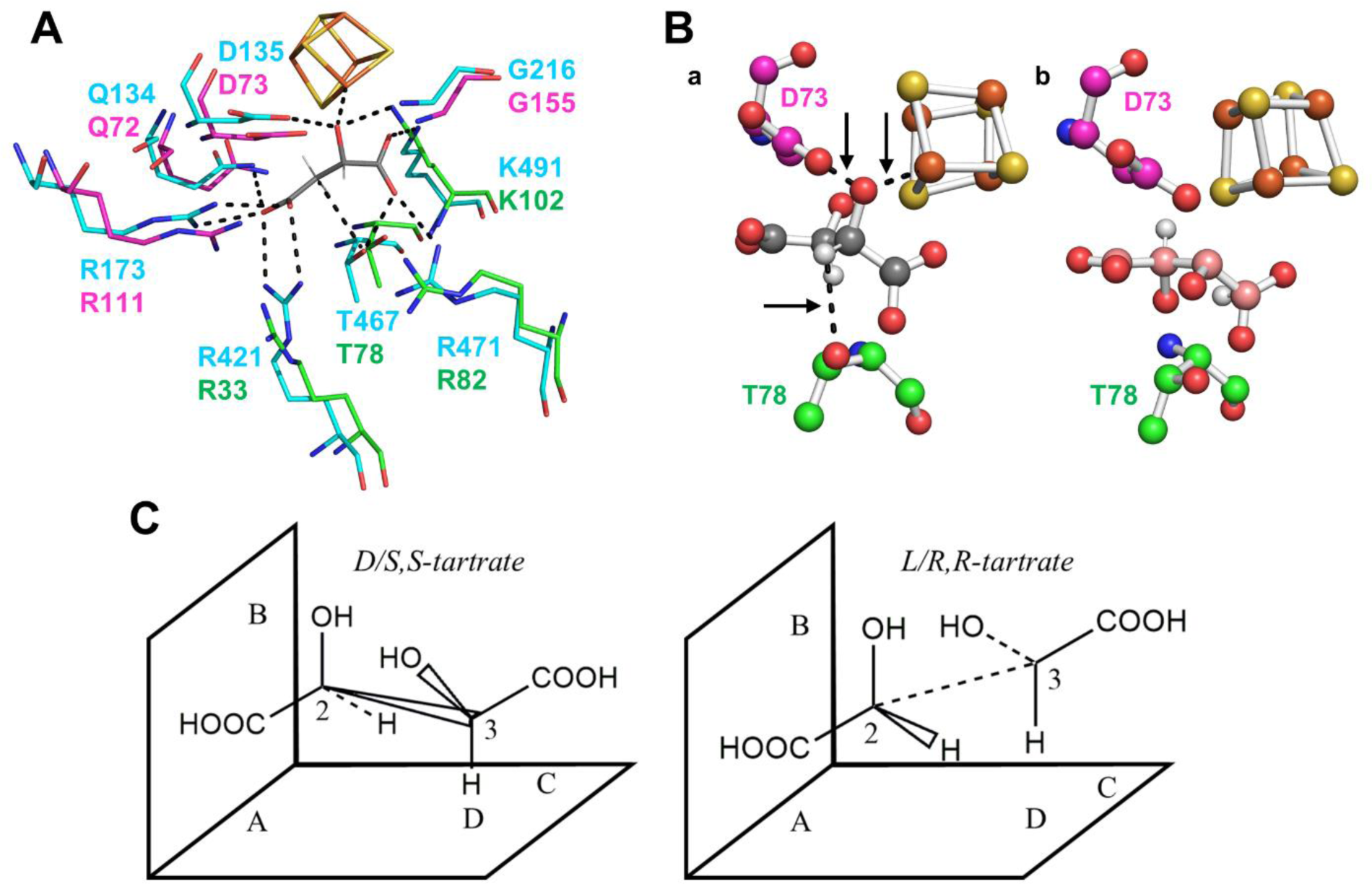
Comparison of the active site in class-I *Lm*FH crystal structure with the modelled structure of *Ec*L-TTD. **A.** Superposition of all the active site residues that hydrogen bond with the substrate (grey) from *Lm*FH structure (cyan) with AlphaFold predicted *Ec*L-TTD structure (α-subunit: magenta; β-subunit: green). Sidechains of R33, R111 and K102 in *Ec*L-TTD have slightly different rotamer conformations while other residues superpose well on *Lm*FH. **B.** Disposition of the Fe-S cluster and the side chains of the catalytic acid, D73 and base T78 in *Ec*L-TTD structure with respect to tartrate. The figure was generated by superposition of the L-malate bound crystal structure of *Lm*FH on to the predicted structure of *Ec*L-TTD. On this superposition, R,R/S,S-tartrate were superposed in the place of the crystallographic substrate R-malate. **a.** C2-OH group of (S,S)-tartrate (grey) contacts the Fe-S cluster and catalytic acid D73,whereas C3-H is oriented in the direction of catalytic base T78 suitable for the catalysis. **b. (**R,R)-tartrate (pink) points in a direction away from both the Fe-S cluster and the catalytic acid D73. The C3-H group also points away from the catalytic base T78 (*Ec*L-TTD numbering). Note that while (R, R)-tartrate is the substrate for *Ec*L-TTD, (S, S)-tartrate does not bind the enzyme. Hence, a reorientation of both the catalytic residues and the substrate is necessary for catalysis to occur. **C.** Schematic of the enantiomer superposition model for the stereospecific recognition of tartrate. A and C are contact points on the enzyme surface which bind the COOH groups. The C2-OH group binds to B that consists of the Fe-S cluster and D73 residue. The fourth contact point D is formed by T78 that binds to C3-H. The flexibility of the C3(OH)COOH group is critical in orienting the fourth contact point in the same direction in both enantiomers.

In class-I FHs, substrate specificity arises out of binding of the C2-OH of the substrate, S-malate with the catalytic aspartate D62 and the Fe-S cluster. However, in the modelled structure of *Ec*L-TTD, the Fe-S cluster, and the catalytic acid D73 (α-subunit) point away from C2-OH of R-malate and are in an orientation not favorable for catalysis (Fig. 4B). As the location of the 3 cysteines binding the 4Fe-4S cluster overlap with that in FH, a reorientation of one or two other critical residues in the active site should lead to switch in stereospecificity. As proposed in the enantiomer superposition model(Bentley, 2003), in the case of *Ec*L-TTD it is the critical flexibility of the C3(OH/H)COOH group which lets R, R-tartrate and R-malate bind in the right orientation to the active site enabling C2(OH) and C3-H to make the correct contacts for the reaction to proceed (Fig. 4C). Indeed, the C3 carbon is responsible for this ‘flexibility’, which orients the C3-H of tartrate, the vital fourth contact point for catalysis, in both the enantiomers in the same direction instead of opposite sides. Hence from the enantiomer superposition model, the distinguishing group i.e., C3-H/OH of tartrate does not have to lie on completely opposite side and a slight reorientation of the active site aspartate and threonine, and the Fe-S cluster along with substrate reorientation and conformer selection about the C2-C3 bond of the dicarboxylic acid should be sufficient for the reaction to proceed on (R, R)-tartrate in *Ec*L-TTD. Similar mechanism has been proposed by Ronald Bentley for isocitrate dehydrogenase for the binding of the diastereomers 2R, 3S and 2S, 3R isocitrate (Bentley, 2003).

To accommodate the chemistry in the active site of L-TTD, we propose a reorientation of the catalytic pocket mediated by the C-terminal residue K201 which makes a direct contact with G155 of the neighboring α subunit, a critical residue of the proximal loop “AGGGC” motif present in the active site (Fig. S5D). This interaction is seen upon superposition of the AlphaFold predicted structures of α- and β-subunits of *Ec*L-TTD on the *Mj*FH crystal structure. *Ec*L-TTD has a C-terminal extension absent in *Mj*FH and the contact of the last residue K201 in the superposed structure with the AGGGC motif could lead to subtle reorganization of the catalytic pocket leading to switch in substrate stereospecificity preference.

## Conclusion

*E. coli* L-tartrate dehydratase and *M. jannaschii* fumarate hydratase share a high degree of sequence conservation and consequently adopt almost identical folds, with very similar disposition of active site functionalities. Yet their favored substrates have opposite chiralities. The phylogenetic proximity of -TTD to archaeal two-subunit FHs suggests that L-TTD may have diverged during evolution to accommodate the binding and catalysis of the opposite enantiomer. As the active site conformation appears identical to that in FH, mutations not directly in the active site must have allowed the switch in stereospecificity. Subtle differences in the active site become evident from our activity measurements on malate and tartrate. In *Mj*FH, inhibition by meso-tartrate and absence of inhibition by R-malate, and activity on S-malate and (S, S)-tartrate, indicate that in this enzyme, the environment around the 4Fe-4S cluster and the α-subunit catalytic acid D62 that contact C2-OH is conformationally rigid while that around the catalytic base T80 in the β subunit that contacts C3 is relatively flexible to accommodate both configurations. Unlike in *Mj*FH, *Ec*L-TTD is active on (R, R)-tartrate and R-malate, and inhibited by both S-malate and meso-tartrate, with (S, S) tartrate showing no effect. This indicates that in *Ec*L-TTD the environment around the 4Fe-4S cluster and the catalytic acid D73 of the α-subunit accommodates both R and S configuration of C2 while the catalytic base T78 of the β-subunit is comparatively fixed accommodating only the R configuration of C3. Despite this, *Ec*L-TTD exhibits strict substrate fidelity and only (R, R) tartrate and R-malate are substrates and differences in the inhibition pattern between *Mj*FH and *Ec*L-TTD only support the presence of subtle differences in active site structure. Clearly, dynamic reorientation of both active site residues and the substrate must occur to rationalize the observed results.

Classical models of enzyme-substrate binding do not explain this unique behaviour, but the enantiomer superposition model provides a possible explanation. A minor reorientation of the active site residues, rather than a complete reversal, may lead to the accommodation of enantiomers/diastereomers in the active site. Conformational plasticity in both the enzyme and the substrate tartrate merit serious consideration. While Koshland’s “four location” hypothesis(Mesecar & Koshland, 2000) may have given way to the “enantiomer superposition model”, recalling his ideas of “induced fit”(Koshland, 1958) may yet provide a basis for discussing the remarkable versatility of enzymes in chiral discrimination. To summarize, the puzzle of stereospecificity in these two well-conserved enzymes highlights the complexity of enzyme-substrate interactions and the enigmatic nature of chiral compounds even after 170 years since their discovery.

## Materials & Methods

### Chemicals, strains, and molecular biology reagents

Restriction enzymes, Phusion polymerase, and T4 DNA ligase were obtained from New England Biolabs, USA. Ni-NTA conjugated agarose beads were obtained from Thermo Fisher Scientific Inc. Malic dehydrogenase from the porcine heart was commercially procured from Sigma-Aldrich (product number M2634), USA. Primers were custom synthesized from Sigma Aldrich, Bengaluru. Media components for growing *E. coli* cultures were from Himedia, Mumbai, India.

### Sequence alignment and phylogenetic tree construction

Sequences of class-I FH and L-TTD were obtained from NCBI BLAST-P tool using *Methanocaldococcus jannaschii* FH (*Mj*FH), *E. coli* FH (*fumA*) and *E. coli* L-TTD protein sequences as queries. Clustal Omega(Goujon et al., 2010; Sievers et al., 2011) was used for multiple sequence alignment and output was processed using ESPRIPT(Robert & Gouet, 2014) for visualization. For the construction of phylogenetic tree, Excel was used to concatenate α and β protein sequences of two-subunit class-I FH and L-TTD. The duplicate sequences were removed and clustered using CD-HIT online suite at a cut-off of 85% sequence identity to remove redundancies. The sequences were then aligned in MEGA X(Kumar, Stecher, Li, Knyaz, & Tamura, 2018) using MUSCLE algorithm. The evolutionary history was inferred by using the Maximum Likelihood method and Le Gascuel model. The tree with the highest log likelihood (-88755.07) is shown. 500 bootstrap replicates(Felsenstein, 1985) were performed and branches corresponding to partitions reproduced in less than 60% bootstrap replicates are collapsed. Initial tree(s) for the heuristic search were obtained automatically by applying Neighbour-Join and BioNJ algorithms to a matrix of pairwise distances estimated using a JTT model, and then selecting the topology with superior log likelihood value. A discrete Gamma distribution was used to model evolutionary rate differences among sites (5 categories (+G, parameter = 1.0890)). The rate variation model allowed for some sites to be evolutionarily invariable ([+I], 2.92% sites). This analysis involved 264 amino acid sequences. All positions with less than 95% site coverage were eliminated, i.e., fewer than 5% alignment gaps, missing data, and ambiguous amino acids were allowed at any position (partial deletion option). There was a total of 428 positions in the final dataset.

### Cloning, protein expression and purification

Generation of expression plasmids carrying genes for *Mj*FH α and β subunits have been described(Bellur et al., 2023). Expression and purification of *Mj*FH was carried out as previously described(Bellur et al., 2023). Genes for the two subunits, L-TTDα and L-TTDβ were PCR amplified using primers listed in Supplementary Table 1 from the genomic DNA of the *E. coli* strain DH5α strain and cloned into two tandem multiple cloning sites of the pET-Duet vector using the enzymes *EcoRI* and *HindIII* for L-TTDα such that the (His)_6_ tag is at its N-terminus and enzymes *NdeI* and *XhoI* for L-TTDβ. Mutations were sequentially incorporated into *Mj*FH using the primers listed in Supplementary Table 1. Four mutants were made; P143M + K144A (**MA**), K144A + S148C (**AC**), P143M + K144A + S148C (**MAC**) and I140D + F142Y + P143M + K144A + S148C + N150L (**DYMACL**). All mutant clones were confirmed by DNA sequencing. All the mutants expressed well and were obtained in good purity.

L-TTD protein was expressed in BL21(DE3)-RIL *E. coli* cells that also carried a second plasmid (RIL) that expressed tRNAs for the rare codons of arginine, isoleucine, and leucine. This combination allowed for the overexpression of the L-TTD protein. BL21 (DE3) RIL *E. coli* cells were transformed with the recombinant plasmid pETduet_LTTD and cultured at 37 °C to an O.D. 600 of ∼1.0 before being induced with 0.3 mM IPTG and grown for an additional 16 hours at 30 °C to overexpress L-TTD α and β subunits. After pelleting, the cells were resuspended in lysis buffer (50mM Tris HCl, pH 8.0, 5% glycerol, 1mM DTT and 1mM PMSF) and lysed using French pressure cell press (Thermo IEC Inc., USA). The lysate was clarified by centrifugation at 30,000 x g for 30 minutes. Without disturbing the pellet, the supernatant was carefully collected, and a 1 mL slurry of Ni-NTA beads (Qiagen) pre-equilibrated with lysis buffer was added. This was done to facilitate the binding of the His-tagged protein. After the 3 hours of binding at 4 °C, the beads were washed with 50 mL of lysis buffer to remove all unbound proteins, and the Ni-NTA-bound protein was then eluted with 200 mM of imidazole in lysis buffer. Pure protein-containing fractions were pooled and dialyzed against a solution containing 50mM Tris HCl, pH 8.0, 5% glycerol, and 1mM DTT. Following dialysis, the fractions were loaded onto a Q-Sepharose anion-exchange column. A linear gradient of increasing NaCl concentration in a buffer containing 50 mM Tris HCl, pH 8.0, 5% glycerol, and 1 mM DTT was used to elute the protein. SDS-PAGE was used to analyze fractions collected at NaCl concentrations greater than 200 mM, pooled according to purity and dialyzed with a buffer containing 50mM Tris HCl, pH 8.0, 5% glycerol and 1mM DTT. The protein concentration was estimated by the Bradford method with bovine serum albumin (BSA) as the standard(Bradford, 1976).

### Reconstitution of Fe-S cluster

The protein obtained after purification and dialysis is the apoprotein lacking the complete 4Fe-4S cluster. Protocol similar to that followed for reconstitution of *Mj*FH(Bellur et al., 2023) was followed for reconstitution of Fe-S cluster into *Ec*L-TTD. Reconstitution was carried out in an anaerobic chamber to prevent oxidation of the cluster. Purified protein was placed inside the chamber and 50x molar excess of DTT was added and stirred for 30 minutes followed by addition of 10x molar excess of each (NH_4_)_2_Fe(SO_4_)_2_ and Na_2_S. The protein solution was stirred for 3 hours to obtain the holoprotein with fully reconstituted iron-sulfur cluster.

### Enzyme activity

All enzyme activity measurements unless specified otherwise, were conducted at 37 ℃ using a water-circulated cell holder fitted to Hitachi U2010 spectrophotometer. Malate to fumarate conversion was monitored as an increase in absorbance at 240 nm (molar extinction coefficient (ε) of fumarate is 2.4 mM^-1^cm^-1^ at 240 nm). The assay mixture contained 1 mM malate in 50 mM Tris-HCl, pH 7.4. Reaction was initiated by the addition of enzyme and monitored spectrophotometrically. Activity on tartrate was monitored using the coupling enzyme, porcine malate dehydrogenase. The reaction was carried out in a final volume of 500 μl in 50 mM Tris-HCl, pH 7.4, containing 250 μM NADH, 2.5 μg MDH with varying concentrations of R,R-tartrate. Conversion of R,R-tartrate to oxaloacetate was coupled to NAD^+^ formation by the coupling enzyme which converts oxaloacetate to R-malate utilizing a molecule of NADH. The reaction was initiated by the addition of 1.44 μg *E. coli* L-TTD and monitored at 340 nm (molar extinction coefficient(ε) of NADH is 6.22 mM^-1^cm^-1^ at 340 nm). Inhibition kinetics of *Mj*FH and *Ec*L-TTD by substrate analogues meso-tartrate, R,R-tartrate and R-malate was examined by fixing the substrate concentration and varying the concentrations of the inhibitors. The data was fit to the equation for competitive inhibition *v*=*V*_max_[S]/(*K*_m_(1+[I]/*K*_i_))+[S] and the *K*_i_ was estimated using nonlinear regression analysis. GraphPad Prism software was used for data fitting and analysis.

### Size-exclusion chromatography

Analytical size-exclusion chromatography was used to examine the co-purified protein’s oligomeric state on an analytical Superdex 200 10/300 GL column (10 mm X 300 mm) (GE Health Care Life Sciences), connected to an AKTA Basic HPLC system with a UV900 detector. The column was equilibrated with Buffer A (50 mM Tris-HCl, pH 8.0, 5% glycerol, 1 mM DTT) and calibrated using molecular weight standards; blue dextran (2000 kDa), β-amylase (200 kDa), alcohol dehydrogenase (150 kDa), bovine serum albumin (66 kDa), and carbonic anhydrase (29 kDa). The void volume of the column matches the elution volume of blue dextran. The ratio of the elution volume to the void volume of these standards was plotted against their log molecular weight to create a standard curve. 100 μl of pure protein (10 mM) that was a complex of α and β subunits was loaded onto the column, and the elution was measured simultaneously at 280 nm and 220 nm. The flow rate was kept at 0.5 ml min^-1^. By extrapolating the elution volume from the standard curve, the complex’s molecular weight was determined.

### Circular dichroism (CD) spectroscopy

All CD spectra were recorded at 25°C using a Jasco J-810 spectrophotometer, at scanning speed of 50 nm/min, and data interval of 1 nm. Protein samples were in 50 mM Tris-HCl pH 8.0, 5% glycerol, and 1 mM DTT at a concentration of 30 μM. Molar ellipticity (*θ_m_*) was calculated using *θ_m_* = m°*M/(10*L*C) where m° is ellipticity in millidegrees; M, molecular weight (g/mol); L, path length (cm) of the cell, and C, concentration in g/L.

### Structural analysis

AlphaFold(Jumper et al., 2021; Varadi et al., 2022) predicted structures of *Ec*L-TTD with the highest confidence levels were obtained from UniProt database, and superposed on the crystal structures of *Mj*FH (PDB ID: 7XKY) and *Lm*FH (PDB ID: 5L2R), and used for analysis.

## Author contribution

AB, conducted experiments; SM, conducted experiments and wrote manuscript; PS, conducted experiments; VJ, conducted experiments, HB, conceptualization, supervision, funding, manuscript writing.

## Acknowledgements

The authors thank P. Balaram for careful reading of the manuscript and critical comments. H.B. acknowledges financial support from the Department of Biotechnology, Ministry of Science and Technology, Government of India Grants BT/PR11294/BRB/10/1291/2014, BT/PR13760/COE/34/42/2015, and BT/INF/22/SP27679/2018; Science and Engineering Research Board (SERB), Department of Science and Technology, Government of India Grant CRG/2019/004150/IBS, EMR/2014/001276; JC Bose Fellowship, SERB; and institutional funding from JNCASR. A.B. was supported by CSIR for junior and senior research fellowships and from the JC Bose fellowship awarded to H.B. V.J. was supported by junior and senior research fellowships from CSIR.

---

**Figure S1.**
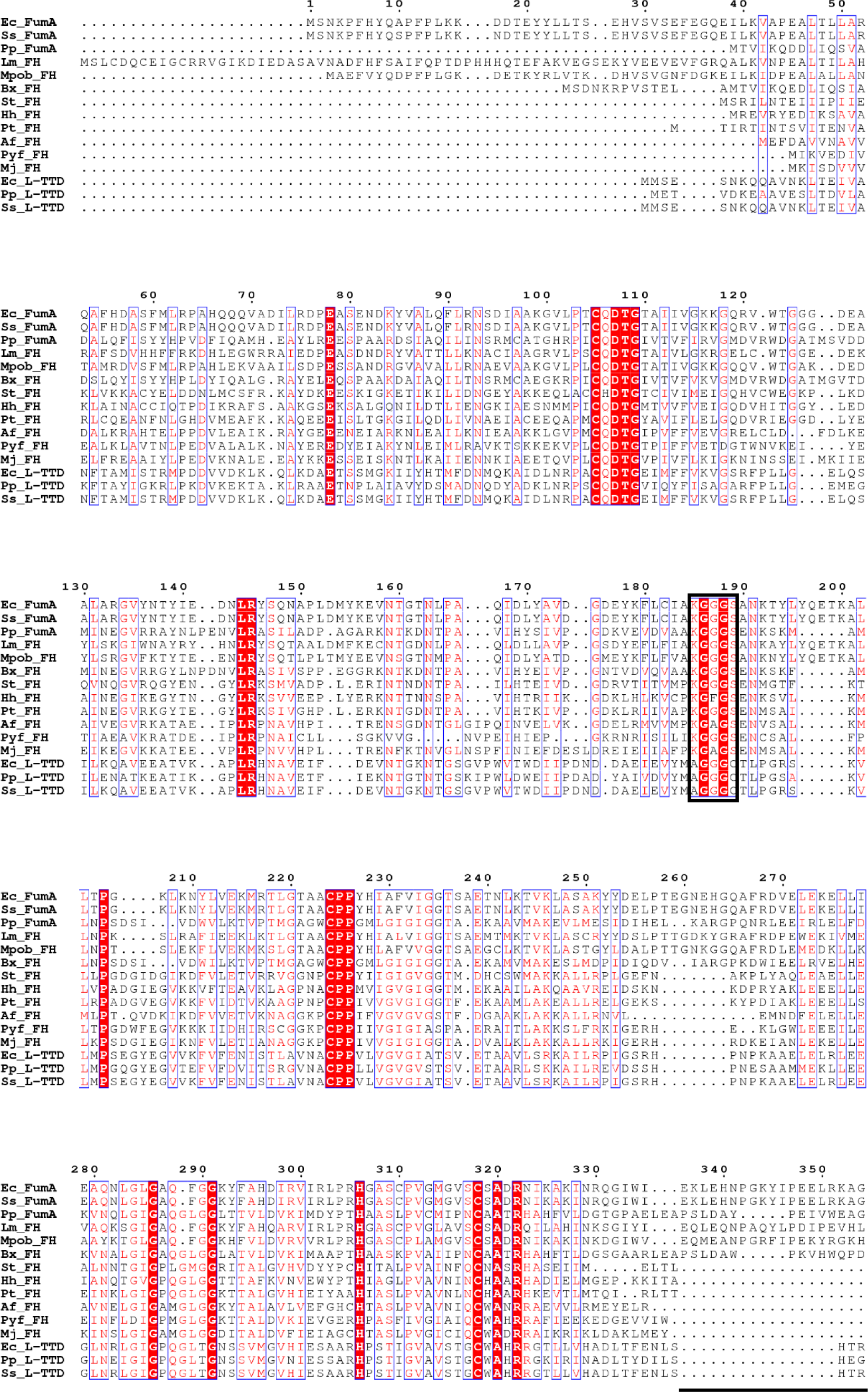
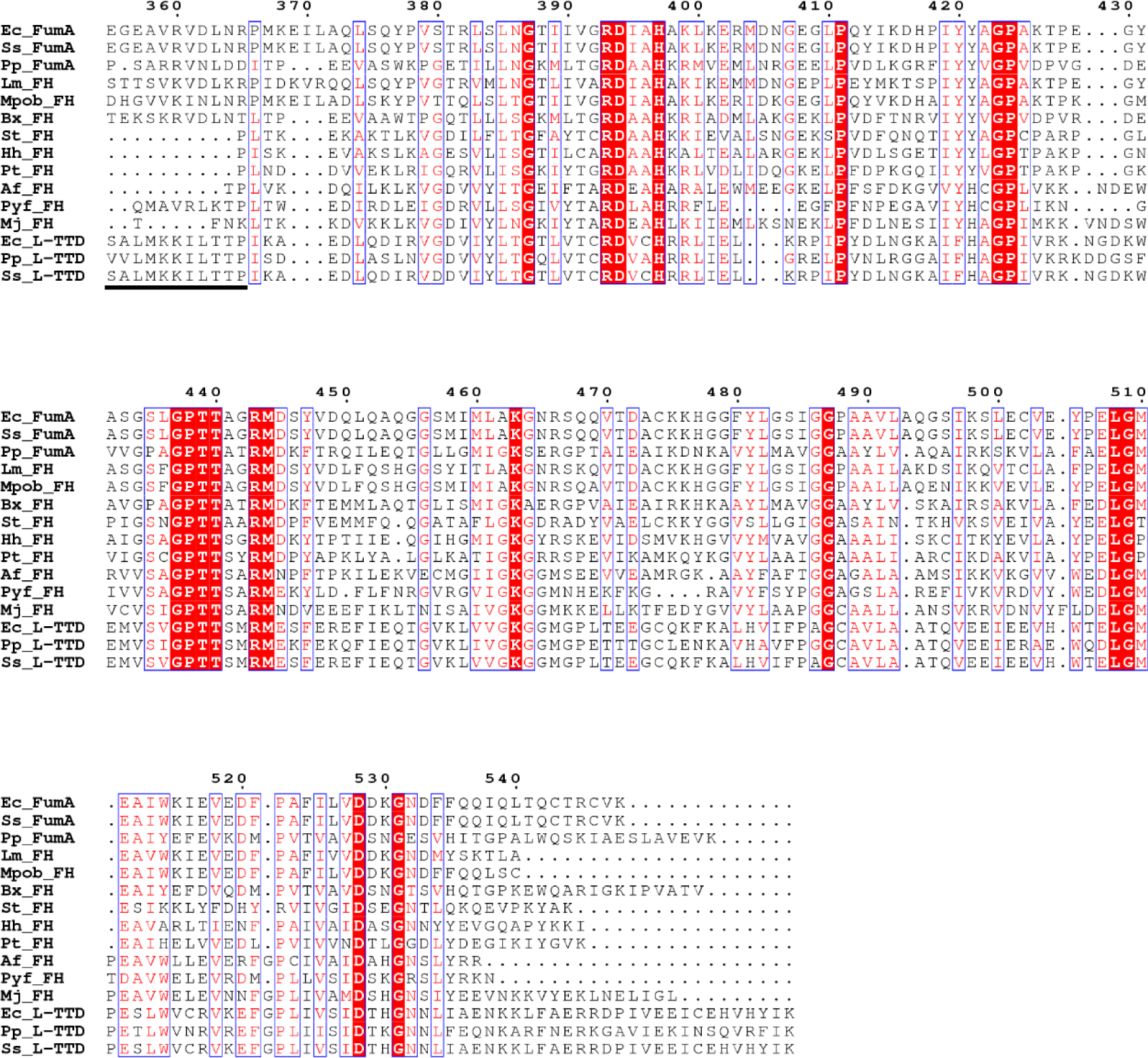
Multiple sequence alignment of biochemically characterized class-I FHs and L-TTDs. KGXGS motif present as an AGGGC motif in L-TTD is enclosed in a box in the alignment. The linker that connects the NTD and CTD of single-subunit FH has been highlighted with a line below the sequences. The α and β subunits of L-TTD and two-subunit FH have been joint as one sequence to generate the alignment. Single subunit FH from *E. coli*, *S. sonnei* and *Pseudomonas putida* are annotated as fumarase A in the alignment. Single subunit FH from Syntrophic propionate-oxidizing bacteria is abbreviated as Mpob in the alignment. FH, fumarate hydratase; FumA, fumarase A; Ec, *Escherichia coli*; Ss, *Shigella sonnei*; Pp, *Pseudomonas putida*; Lm, *Leishmania major*; Mpob, Syntrophic propionate-oxidizing bacteria; Bx, *Burkholderia xenovorans*; St, *Salmonella typhimurium*; Hh, *Helicobacter hepaticus*; Pt, *Pelotomacum thermopropionicum*; Af, *Archaeoglobus fulgidus*; Pyf, *Pyrococcus furiosus*; Mj, *Methanocaldococcus jannaschii*.

**Figure S2.**
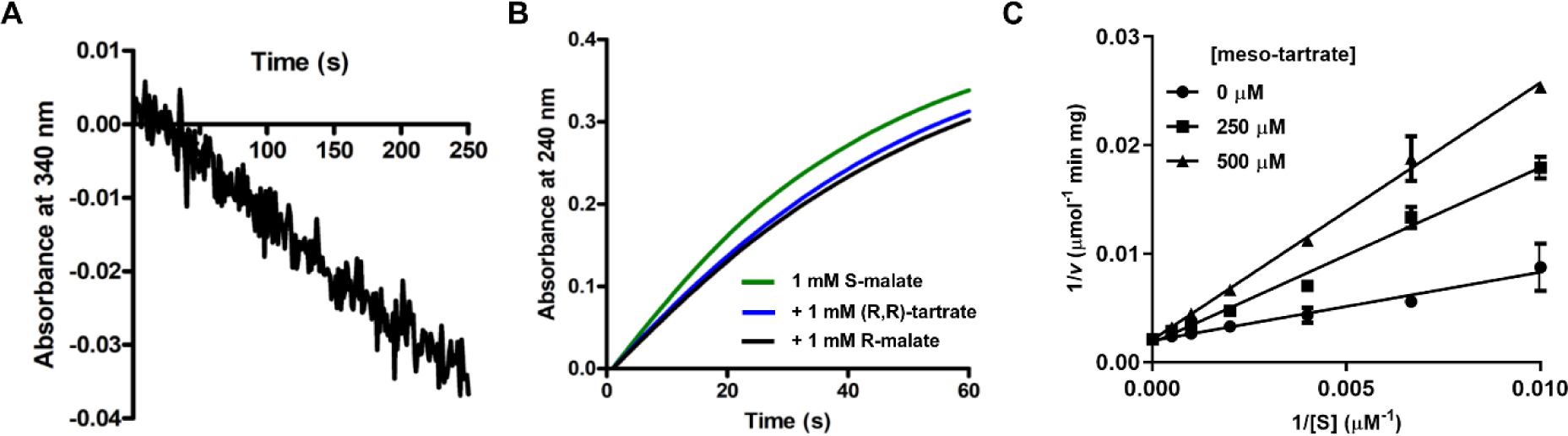
Substrate specificity of *Mj*FH. **A.** Representative progress curve for the conversion of (S, S)-tartrate to oxaloacetate by *Mj*FH. It should be noted that *Mj*FH is a thermostable enzyme with significantly increased activity at 70 ℃. As the coupling enzyme, porcine malate dehydrogenase is optimally active at 37 ℃, the assay was performed at this lower temperature. Hence, the lower activity of *Mj*FH on (S, S)-tartrate as compared to its activity on S-malate, which involves a direct measurement at 240 nm with the assay temperature being 70 ℃. **B.** Representative progress curves for the conversion of 1mM S-malate to fumarate by *Mj*FH (green line) in the presence of 1mM (R, R)-tartrate (blue) and 1 mM R-malate (black) going to show that *Mj*FH is not inhibited by either (R, R)-tartrate or R-malate. **C.** Double reciprocal plots with meso-tartrate as inhibitor and S-malate as substrate. Concentration of S-malate was varied at different fixed concentrations of meso-tartrate (0, 250 and 500 μM). All experiments were repeated twice to confirm reproducibility.

**Figure S3.**
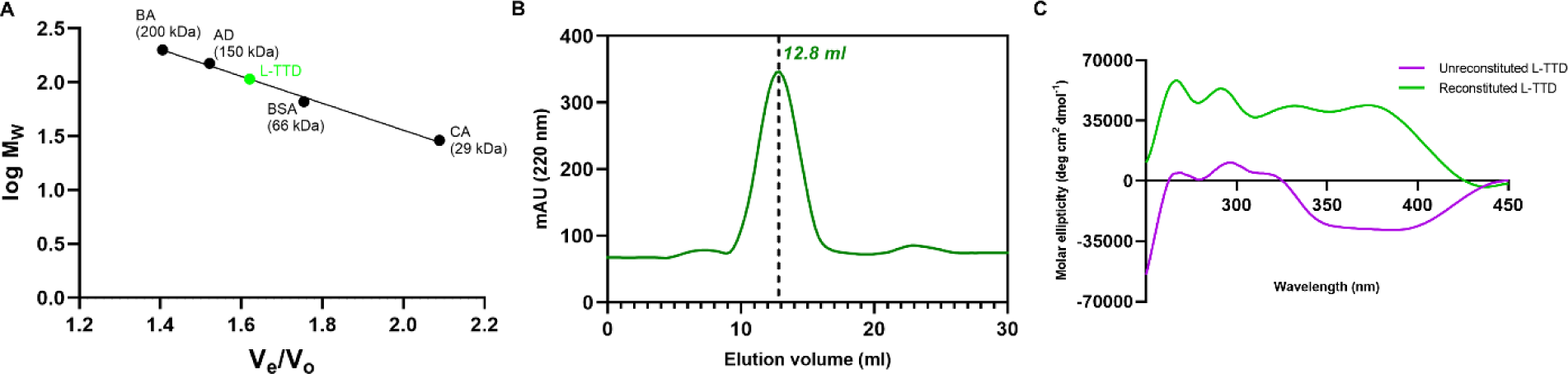
Biochemical characterization of *Ec*L-TTD. **A.** Molecular mass of *Ec*L-TTDαβ complex. The interpolation of the elution volume on the standard curve yields a mass of 107.15 kDa. The standard curve was generated using the protein standards, ADH, alcohol dehydrogenase; BA, β-amylase; BSA, bovine serum albumin; CA, carbonic anhydrase. The elution volume of the standards is indicated with a black filled circle while that of the L-TTDαβ complex by a green filled circle. The elution volume of the complex corresponds to a dimer of heterodimer (2α + 2β). **B.** The complex elutes as a single peak on size-exclusion chromatography. **C.** CD spectra of Fe-S reconstituted and unreconstituted *Ec*L-TTD. Unlike the unreconstituted protein, the reconstituted protein exhibits a positive Cotton effect. All experiments were repeated twice to confirm reproducibility.

**Figure S4.**
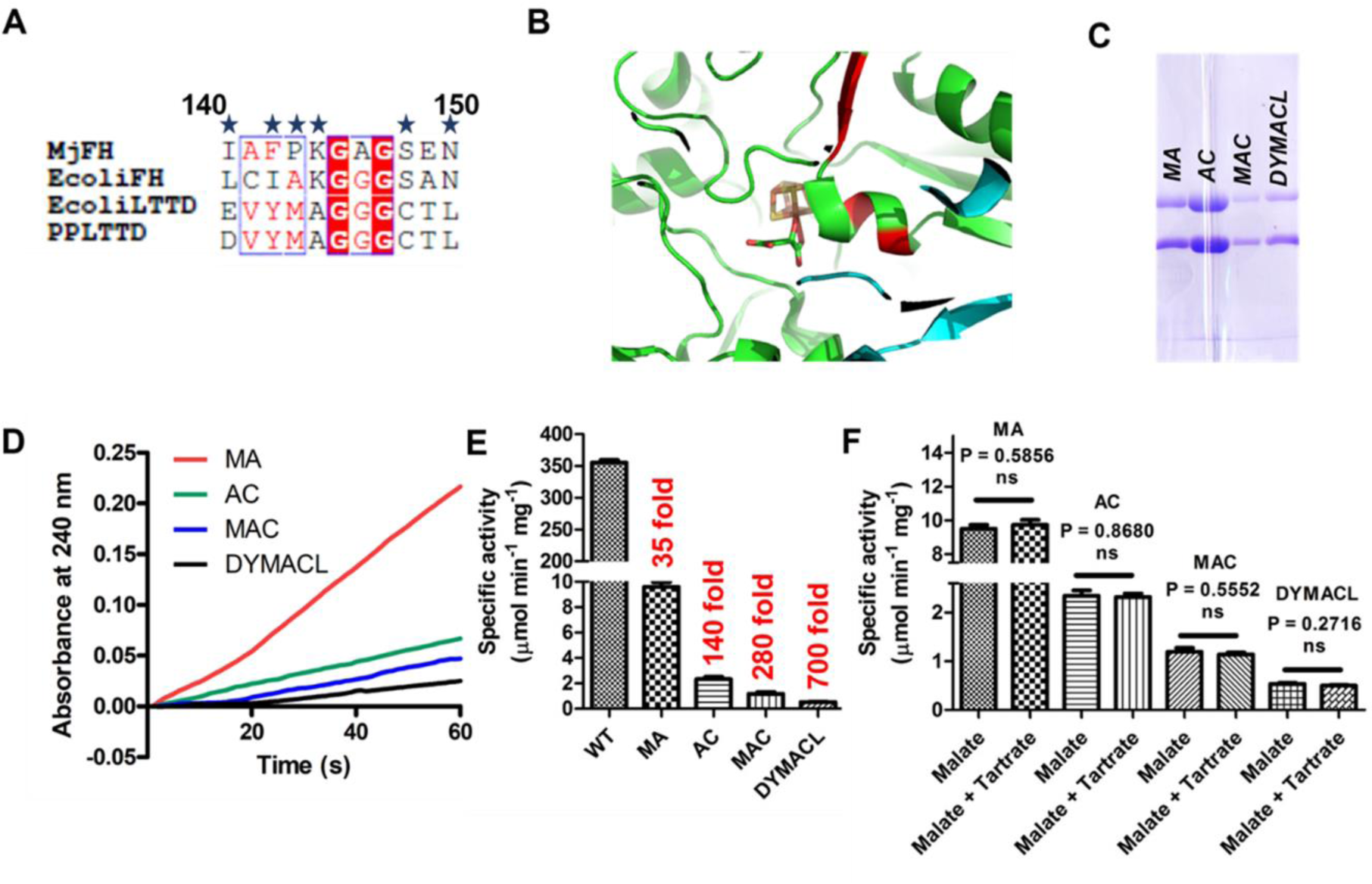
Swapping the residues in the conserved motif from *Ec*L-TTD onto *Mj*FH does not lead to switch in stereospecificity. **A.** Sequence alignment of class-I FHs and L-TTDs shows that residues at positions 140, 142, 143, 144, 148 and 150 are conserved differently in class-I FH and L-TTD. **B.** Residues that correspond to the above 5 positions are located in a helix and a sheet close to the bound substrate (S-malate) and Fe-S cluster in the LmFH structure (5l2r) and are highlighted in red. **C.** SDS-PAGE showing the purified *Mj*FH mutants. Lane 1, MA; lane 2, AC; lane 3, MAC; and lane 4, DYMACL. Abbreviations used to denote the mutants are as follows: P143M + K144A (**MA**), K144A + S148C (**AC**), P143M + K144A + S148C (**MAC**) and I140D + F142Y + P143M + K144A + S148C + N150L (**DYMACL**) **D.** Representative progress curves for conversion of 1mM L-malate to fumarate by mutants MA (red line), AC (green line), MAC (blue line) and DYMACL (black line). **E.** Bar graph for the specific activity of mutants MA, AC, MAC and DYMACL with 1 mM S-malate as substrate showing the fold change in activity in comparison to WT *Mj*FH. **F.** Bar graph showing the comparison of specific activity for the mutants MA, AC, MAC and DYMACL with 1 mM S-malate as substrate in the presence/absence of 1 mM (R, R)-tartrate in the assay mixture. Statistical analysis was done using Student’s unpaired t-test using GraphPad Prism.

**Figure S5.**
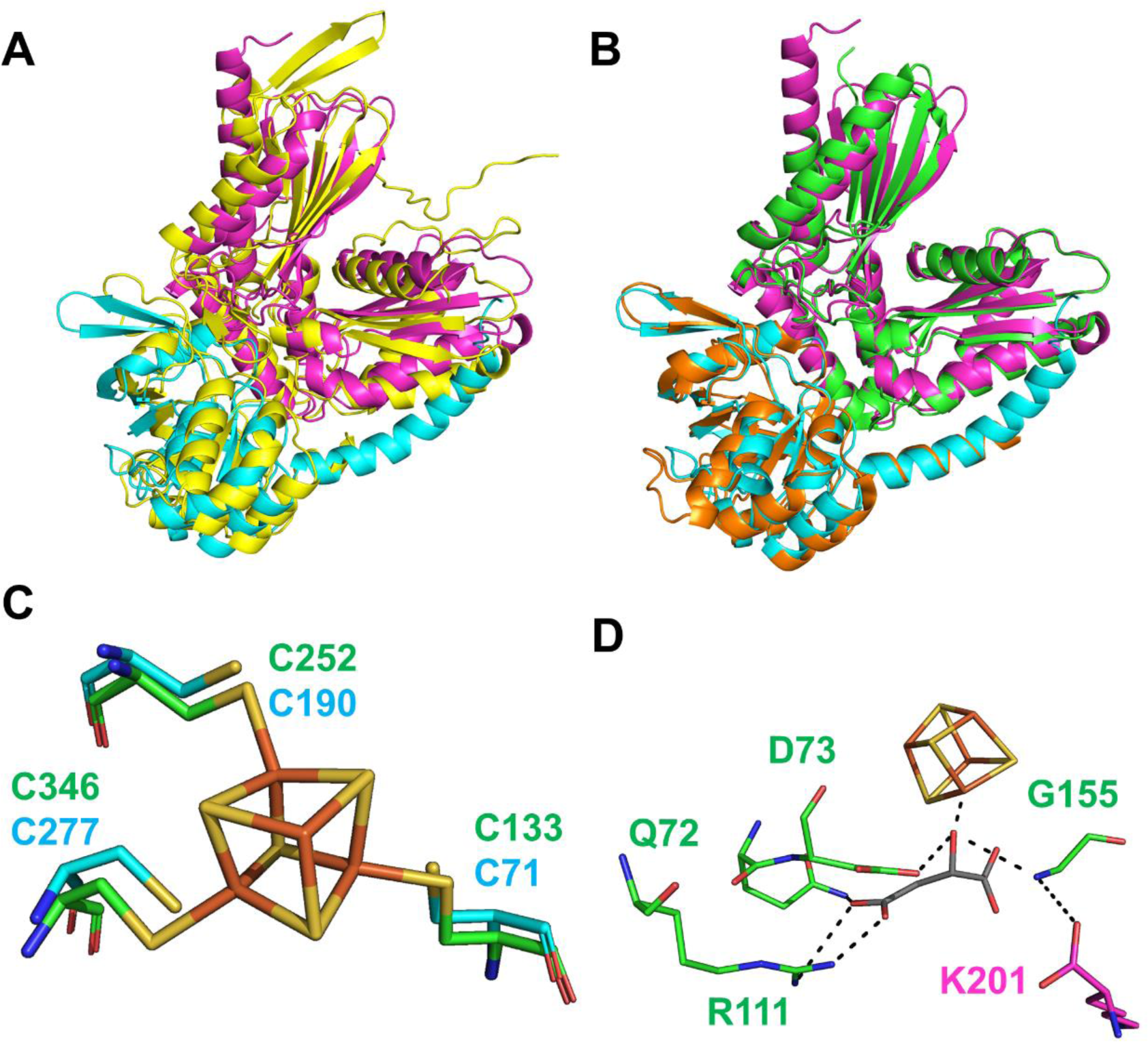
Superposition of AlphaFold predicted structures of *Ec*L-TTD on class-I FH crystal structures. **A.** Superposition of AlphaFold predicted *Ec*L-TTDα (magenta) and *Ec*L-TTDβ (cyan) structures over the *Lm*FH structure (yellow). The two structures align well with RMSD of 1.95 Å, and 1.52 Å, respectively. **B.** Superposition of AlphaFold predicted *Ec*L-TTDα (magenta) and *Ec*L-TTDβ (cyan) structures over the *Mj*FHαβ structure (α: green, β: orange). The two structures align well with a RMSD of 1.104 Å and 1.008 Å, respectively. **C.** Superposition of cysteine residues (C71, C190 and C277) that ligate Fe-S cluster in *Ec*L-TTD predicted structure over cysteine residues (C133, C252 and C346) in *Lm*FH structure. Fe-S cluster from *Lm*FH structure is retained in the superposed snapshot. **D.** *Ec*L-TTD tetramer aligned on *Mj*FH structure reveals that the C-terminal K201 of the β subunit makes contacts with G155 of the “AGGGC” motif of the α subunit in the adjacent hetero dimer of the tetramer. It should be noted that this G155 contacts the C2-OH group of the substrate and the catalytic acid D73.

**Supplementary Table 1.**
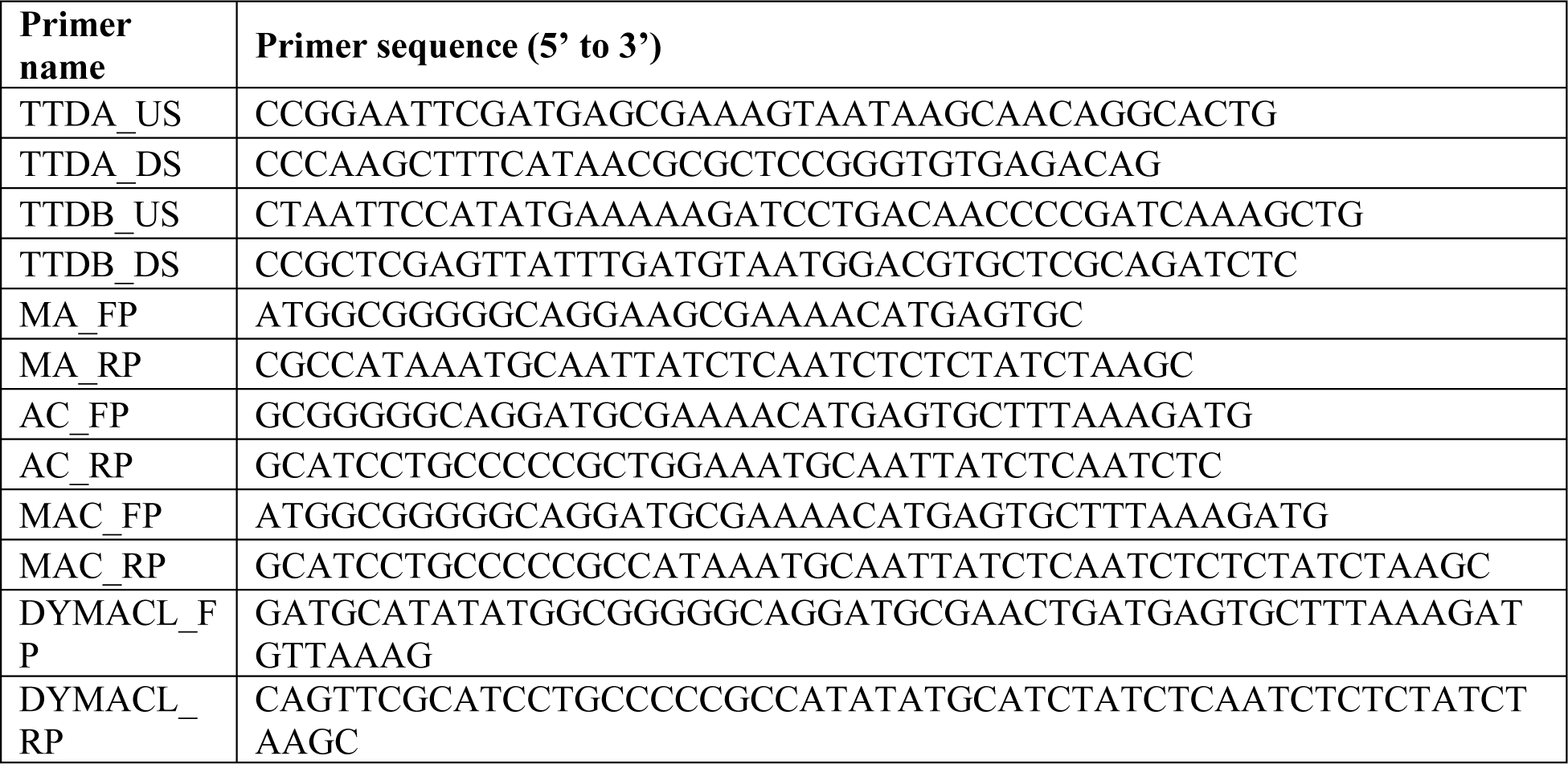
List of primers.

## Notes

### Competing Interest Statement

The authors have declared no competing interest.

